# Identification of a new *Plasmodium falciparum* E2 ubiquitin conjugating enzyme

**DOI:** 10.1101/2024.10.06.616869

**Authors:** Cameron Smith, Mohsen Hajisadeghian, Gerbrand J. van der Heden van Noort, Adán Pinto-Fernández, Benedikt M Kessler, Katerina Artavanis-Tsakonas

## Abstract

The ubiquitin-proteasome system (UPS) is essential for Plasmodium falciparum survival and represents a potential target for antimalarial therapies. We utilised a ubiquitin-activity based probe (Ub-Dha) to capture active components of the ubiquitin conjugating machinery during asexual blood-stage development. Several E2 ubiquitin-conjugating enzymes, the E1 activating enzyme, and the HECT E3 ligase PfHEUL were identified and validated through in vitro ubiquitination assays. We also demonstrate selective functional interactions between PfHEUL and a subset of both human and *P. falciparum* E2s. Additionally, the Ub-Dha probe captured an uncharacterized protein, PF3D7_0811400 (C0H4U0) with no known homology to ubiquitin-pathway enzymes in other organisms. Through structural and biochemical analysis, we validate it as a novel E2 enzyme, capable of binding ubiquitin in a cysteine-specific manner. These findings contribute to our understanding of the *P. falciparum* UPS, identifying promising novel drug targets and highlighting the evolutionary uniqueness of the Ub-proteasome system in this parasite.

## Introduction

Ubiquitin is a small (8.6 kDa) post-translational protein modifier that, when conjugated to proteins, modulates their activity, fate, and localisation within the cell. The machinery that mediates ubiquitination and deubiquitination of substrates is complex, highly regulated, and functionally conserved across eukaryotes. During the blood-stage of malaria infection, the *Plasmodium* parasite relies on ubiquitin-driven protein turnover to progress through its life cycle. This process ensures the timely removal of stage-specific proteins, which is essential for parasite development and survival.. Additionally, ubiquitination and subsequent degradation of misfolded proteins have been shown to maintain homeostasis during heat shock induced by host fever spikes ^1^). More recently, this modification has garnered increased attention due to its involvement in frontline drug resistance. Artemisinin causes oxidative damage to parasite proteins whose clearance is dependent on the ubiquitin-proteasome system. As such, interference with the ubiquitin pathway by ways of proteasome inhibition, augments artemisinin toxicity. Exploiting this synergy could be particularly useful in the context of resistance whereby co-administering a ubiquitin pathway inhibitor could reestablish the potency of artemisinin ^2^.

The groups of enzymes which constitute the ubiquitin proteasome system (UPS) possess specific conserved domain motifs and architectures that facilitate their function and allow for *in silico* identification from genomic sequences. Despite the substantial divergence between Apicomplexa and model eukaryotic organisms, there are recognisable homologs to all groups of ubiquitin-proteasome system (UPS) enzymes in *Plasmodium*, i.e. E1 activating, E2 conjugating, E3 ligating and deubiquitinating (DUB) enzymes. Two activating enzymes have been identified in *P. falciparum* parasites, PF3D7_1225800 is found in the cytoplasm and PF3D7_1333200 in the apicoplast ^3,4^. In addition, there are 12 putative E2 ubiquitin conjugating enzymes, 4 HECT E3 ligases, 2 types of cullin-RING ligases, 42 putative RING family E3 ligases, and 28 putative deubiquitinating enzymes ^5^. Yet, there are approximately 1,500 proteins with unknown function and domain composition in the *Plasmodium* proteome and several hundred million years of evolutionary divergence between *Plasmodium* and model organisms. Therefore, there are likely to be highly divergent and novel enzymes with roles in the *Plasmodium* UPS that have not yet been identified.

Within the UPS, the points of entry (E1 ubiquitin activating enzyme) and exit (the proteasome and DUB enzymes) have been experimentally demonstrated to be essential for parasite survival ^3,6,7^. However, the enzymes and processes between these events that confer specificity and regulation to this pathway through differential substrate recognition and ubiquitin chain formation have, to date, been largely overlooked as potential therapeutic targets. Given greater sequence and likely functional divergence of E2 and E3 enzymes between *Plasmodium* and the human host, the identification and characterisation of these enzymes would provide novel targets and enable species-specific therapeutic intervention.

Activity-based probes (ABPs) are biochemical tools that have been developed to delineate component interactions and interdependencies within the UPS, allowing insight into the spatiotemporal organisation and dynamics of the ubiquitin network. ABPs consist of three parts, each performing a different function: a reactive group whose chemistry allows for covalent bond formation with the desired targets’ active sites in a physiological environment; a recognition element that drives the protein-protein interaction specificity with the desired target; and a reporter tag that allows for identification of the ABP-substrate conjugate ^8^.

The Ub-Dha probe demonstrates specificity to ubiquitination machinery through its requirement for ATP-mediated activation by the E1 activating enzyme ^9^. It achieves ubiquitination enzyme capture by either reacting natively through its C-terminal carboxyl group with the active site cysteine of a ubiquitination enzyme, or through its dehydroalanine (Dha) group which results in the formation of an irreversible thioether-linked adduct. The stochasticity of these reactions results in a cascade whereby either the Ub moiety can transverse the ubiquitination pathway by successive transthiolation reactions or at any given step can covalently trap an enzyme by 1,4-addition.

Here, we interrogate the active ubiquitin machinery of asexual, erythrocytic stage *Plasmodium falciparum* using a Ub-Dha activity-based probe (ABP) for protein capture, enrichment, and identification by mass spectrometry. We identify the *P. falciparum* E1, several E2 enzymes, a HECT E3 ligase, and a highly enriched unknown protein. We validate these enzymes as bona fide UPS components and determine which E2s serve as functional partners of the identified HECT, PfHEUL, an enzyme that has been implicated in parasite virulence in mouse model systems ^10^. We also characterise the unknown protein as a novel E2 enzyme.

## Materials and Methods

### Asexual parasite stage culture

*Plasmodium falciparum* (strain NF54) was cultured essentially as described in ^11^. Briefly, parasites were cultured within O+ human erythrocytes provided by anonymous health donors, via the National Health Service (NHS) Blood and Transplant, at 5% haematocrit in RPMI 1640 complete medium containing L-glutamine and 25 mM HEPES (Sigma), supplemented with 0.2% sodium bicarbonate, 0.5% Albumax II (Gibco), 0.36 mM hypoxanthine (Sigma) and 50 mg/L gentamicin sulphate (Melford) under a 90% N2/5% O2/5% CO2 gaseous atmosphere at 37°C, within a hypoxic incubator chamber (Billups-Rothenberg). Parasites were maintained at a parasitaemia of 0.2-10%, calculated by counting the percentage of parasitized erythrocytes of Hemacolor (Sigma) stained thin blood smears using light microscopy.

Infected erythrocytes at a parasitaemia of 8-10% were collected by centrifugation, and washed twice in 10 v:v PBS. A volume of 0.2% (w/v) saponin equivalent to the starting culture was used to resuspend the cell pellet, and incubated for 10 minutes at 4°C. Following centrifugation at 3,200 g, the supernatant was removed, and the parasite pellet was washed three times in ice cold PBS. Following the final wash, the supernatant was aspirated, and the pellet flash-frozen in liquid nitrogen and stored at -80°C until use.

### Methods for Ubiquitin-Dha activity-based protein profiling (ABPP)

1 L of P.falciparum culture at 8 % parasitaemia was grown and harvested. Frozen parasite pellets were resuspended in lysis buffer (50 mM HEPES pH 8, 100 mM NaCl, 1 mM DTT, 1% w/v N-octyl glucoside, Complete Mini protease inhibitor). The cell suspension was lysed by 5 freeze-thaw cycles with agitation. The soluble fraction was separated by centrifugation, and protein concentration measured by the Pierce BCA protein assay kit (ThermoFisher) following the manufacturer’s instructions.

The procedure for the biotin-Ub-Dha probe used was based on the method described in ^9^. For use with the biotin-Ub-Dha probe, the supernatant was split in two (experimental and control) and 10 mg total protein was used in each condition. To the control sample, 2 units of apyrase (Sigma-Aldrich) were incubated with the lysate for 30 minutes at 37°C to deplete residual ATP. 10 μg of biotin-Ub-Dha was added to both lysates, and 10 mM Mg-ATP was added only to the experimental lysate. Both lysates were subsequently incubated for 90 minutes at 37°C with agitation; within this time the experimental lysate was supplemented with 1 mM Mg-ATP every 15 minutes.

Following incubation, 20 μL pre-washed Pierce Neutravidin resin (ThermoFisher) was added and incubated for 3 hours at 4°C with rotation. Beads were collected by centrifugation and washed 4 times with a change of vessel before the final wash. (Wash 1: 2% SDS. Wash 2: 50 mM HEPES pH 7.5, 1 mM EDTA, 500 mM NaCl, 1% Triton-X 100, 0.1% sodium deoxycholate. Wash 3: 10 mM Tris pH 8, 1 mM EDTA, 0.5% NP 40, 250 mM LiCl. Wash 4: 50 mM Tris pH 8, 50 mM NaCl). The beads were collected by centrifugation and resuspended in a 1x LDS buffer (ThermoFisher) before incubation at 95°C for 10 minutes. The eluent was collected and processed for LC-MS/MS.

### Proteomics sample preparation

The eluates were prepared for mass spectrometry (MS) analysis following a protocol as described previously ^12^. Briefly, immunoprecipitated sample eluates were diluted to 175 μL with ultra-pure water and reduced with 5 μL of DTT (200 mM in 0.1 M Tris, pH 7.8) for 30 minutes at 37°C. The samples were then alkylated with 20 μL of iodoacetamide (100 mM in 0.1 M Tris, pH 7.8) for 15 minutes at room temperature, protected from light.

Protein precipitation was performed using a double methanol/chloroform extraction method. Protein samples were treated with 600 μL of methanol, 150 μL of chloroform, and 450 μL of water, followed by vigorous vortexing. Samples were centrifuged at 17,000 × g for 3 minutes, and the upper aqueous phase was removed. Proteins were pelleted by adding 450 μL of methanol and centrifuging at 17,000 × g for 6 minutes. The supernatant was discarded, and the extraction process was repeated. The precipitated proteins were resuspended in 50 μL of 6 M urea and diluted to less than 1 M urea with 250 μL of 20 mM HEPES (pH 8.0) buffer.

Protein digestion was conducted by adding trypsin (Worthington; from a 1 mg/mL stock in 1 mM HCl) at a ratio of 1:100, rocking at 12 rpm at room temperature overnight. Following digestion, the samples were acidified to 1% trifluoroacetic acid, desalted on C18 solid-phase extraction cartridges (SEP-PAK plus, Waters), dried, and re-suspended in buffer A.

Liquid chromatography-tandem mass spectrometry (LC-MS/MS) analysis was performed using a Dionex Ultimate 3000 nano-ultra high-pressure reverse-phase chromatography system coupled online to a Q Exactive (Thermo Scientific) mass spectrometer, as described previously ^13^. In brief, the samples were separated on an EASY-Spray PepMap RSLC C18 column (500 mm × 75 μm, 2 μm particle size, Thermo Scientific) over a 60-minute gradient (120 minutes for the matching proteome) of 2–35% acetonitrile in 5% dimethyl sulfoxide (DMSO), 0.1% formic acid at a flow rate of 250 nL/min. MS1 scans were acquired at a resolution of 60,000 at 200 m/z, and the top 12 most abundant precursor ions were selected for high collision dissociation (HCD) fragmentation.

### Data Analysis

Raw MS data was used as input for analysis using MaxQuant software v.2.0.3.1. Parameters were set as default with exceptions: deamidation (NQ) and GlyGly (K) were included as variable modifications; label min ratio count was set to 1 and deamidation was included as a modification used in protein quantification, while discard unmodified counterpart peptides was unchecked; the FTMS MS/MS was set to 0.05 Da and the ITMS MS/MS match tolerance was set to 0.6 Da; and iBAQ LFQ was checked. The PlasmoDB v61 proteome fasta file was used as the reference proteome file.

The resultant ProteinGroup output file was input into Perseus v1.6.15.0 for further analysis using the iBAQ LFQ values. Column values were filtered based on categorical column: only identified by site, reverse, and potential contaminant. Samples were grouped as control or experimental by categorical row annotation. iBAQ LFQ values were transformed by log2(x), and invalid values filtered with a minimum requirement of 2 valid values per group in the case triplicates. Invalid values were imputed from a normal distribution: width (0.3), down shift (1.8). A scatter plot was produced using fold change as the x-axis and log10(average intensity) as the y-axis, both calculated from the LFQ iBAQ. A significance B outlier test was performed on this plot to identify statistically enriched hit proteins at a threshold p<0.05.

### Protein expression and purification

pET28a+ plasmids encoding His-tagged *Plasmodium falciparum* proteins were heat shocked into C41 *E.coli* and selected by kanamycin. 2xYT media was inoculated with a transformed colony, and protein expression induced with IPTG at 0.8 OD600. Following overnight growth, culture was harvested by centrifugation, and mechanically lysed in 50 mM Tris pH 8, 500 mM NaCl, 0.5 mM TCEP supplemented with Complete protease inhibitor tablets and benzonase, using a cell disruptor (Constant Systems). Soluble protein was taken following clarification by centrifugation and 0.22um membrane. Proteins were subject to Ni-affinity by HisTrap Excel column and size exclusion chromatography by either S75 16/60 or S200 10/300 column, along with an AKTA system (Cytiva). A desalting step was incorporated into the workflow at the point of elution from the IMAC resin, such that upon imidazole-mediated elution the protein was minimally exposed to the high salt concentration of the elution buffer. Eluted fractions were subject to analysis by SDS-PAGE and Coomassie staining, before aliquoting and storage at -80°C following flash-freezing. For PfHEUL, protein expression was achieved using optimised conditions based on expression screening - 3 hours at 37°C following induction with 1 mM IPTG. *E.coli* cell lysate containing the recombinant PfHEUL HECT domain was subject to nickel-based immobilised metal affinity chromatography (IMAC).

### SDS-PAGE and Western blot

SDS-PAGE was performed using NuPAGE 4-12% Bis-Tris precast gels using MOPS or MES buffer (ThermoFisher). a 4x LDS buffer containing 5% β-mercaptoethanol was added to samples in a 1:3 ratio, this mixture was then heated to 95°C for 10 minutes prior to gel loading. The gel was run at 180V for 40-60 minutes. Coomassie staining of SDS-PAGE gels was performed using InstantBlue gel stain following the manufacturer’s instructions (Expedion).

SDS-PAGE gels were transferred to methanol-activated PVDF membrane by Trans-Blot SD semi-dry transfer at a constant voltage of 10V for 1 hour (BioRad). Membranes were washed with TBST before blocking in 5% milk (TBST) for 1 hour with agitation. Primary (and conjugated) antibodies: rat anti-HA-peroxidase clone 3F10 high affinity (Roche), mouse anti-His clone 1B7G5 (Proteintech), mouse anti-Ub VU-1 (LifeSensors), or mouse anti-PfHEUL HECT (kindly provided by Andrew Blagborough) were incubated with the membrane at a 1:1000 dilution (5% milk-TBST) for 1 hour at room temperature. Following incubation, membranes were washed 3 times with TBST. For the unconjugated antibody, the membrane was incubated again in 5% milk (TBST) containing a 1:5000 dilution of secondary antibody goat anti-mouse-HRP (ThermoFisher) for 1 hour at room temperature. Following antibody incubations the membranes were washed three times in TBST. Probed membranes were visualised using ECL western blotting substrate (Promega) and Azure c500 gel imaging systems.

### Ubiquitination assays

To test the auto-ubiquitination activity of the PfE2 enzymes, 5 uM of *P. falciparum* protein (PF3D7_0527100, PF3D7_0921000, PF3D7_1033900, PF3D7_1203900, PF3D7_1356300) was incubated with 25 uM HA-Ub and 100 nM HsUBA1 (E1), 10 mM ATP, and the reaction buffer components of the Ubiquitin conjugation initiation kit (K-995, R&D systems) following the manufacturer’s instructions.

To validate the autoubiquitination activity of the WT and C8558A PfHEUL HECT domain, 5 uM of the recombinant HECT domain were incubated with 2.5 uM human E2 ubiquitin conjugating enzymes (UBE2D1, UBE2D2, UBE2D3, UBEE21, UBE2L3 from the UbcH (E2) Enzyme kit, K-980D, R&D systems), 100 nM HsUBA1 (E1) and 10 mM ATP in ubiquitin conjugation reaction buffer (R&D systems).

To assess the cooperativity of the *P.falciparum* E1, E2, and HECT E3 domain activities, 100 nM of PfE1 (PF3D7_1225800), 5 uM PfE2 enzymes (PF3D7_0527100, PF3D7_0921000, PF3D7_1033900, PF3D7_1203900, PF3D7_1356300) , and 5 uM of PfHEUL (PF3D7_0826100) HECT domain were incubated with 10 mM ATP in ubiquitin conjugation reaction buffer (R&D systems). All reactions were incubated at 37C for 1 hour. The reactions were quenched with 1X LDS sample buffer (ThermoFisher) supplemented with 50 mM DTT. Samples were analysed by SDS-PAGE and western blot as described above.

### In-silico Analysis

Sequence alignment and residue conservation were conducted using Clustal Omega^14^ and visualised with ESPript 3^15^. Protein 3D structure predictions and protein-protein interaction modelling were carried out using AlphaFold3, a deep-learning-based method^16^. Structural figures were generated using UCSF ChimeraX^17^, and structural alignments were performed with the RCSB PDB Pairwise Structure Alignment tool^18^. To identify orthologs, the Consurf web server^19^ was employed, which also colour-coded the input PDB file based on the computed conservation score for each amino acid residue.

## Results

### A cascading Ub-derivatised probe captures Ub conjugation machinery in *P. falciparum*

The Ub-Dha probe demonstrates specificity to ubiquitination enzymes through its requirement for ATP-mediated activation by the E1 activating enzyme ^9^. It achieves ubiquitination enzyme capture by either reacting natively through its C-terminal carboxyl group with the active site cysteine of a ubiquitination enzyme, or through its dehydroalanine group which forms an irreversible thioether-linked adduct. The stochasticity of these reactions results in a cascade whereby either the Ub moiety can transverse the ubiquitination machinery by successive transthiolation reactions or at any given step can covalently trap an enzyme by 1,4-addition (Figure 1A).

**Figure 1.**
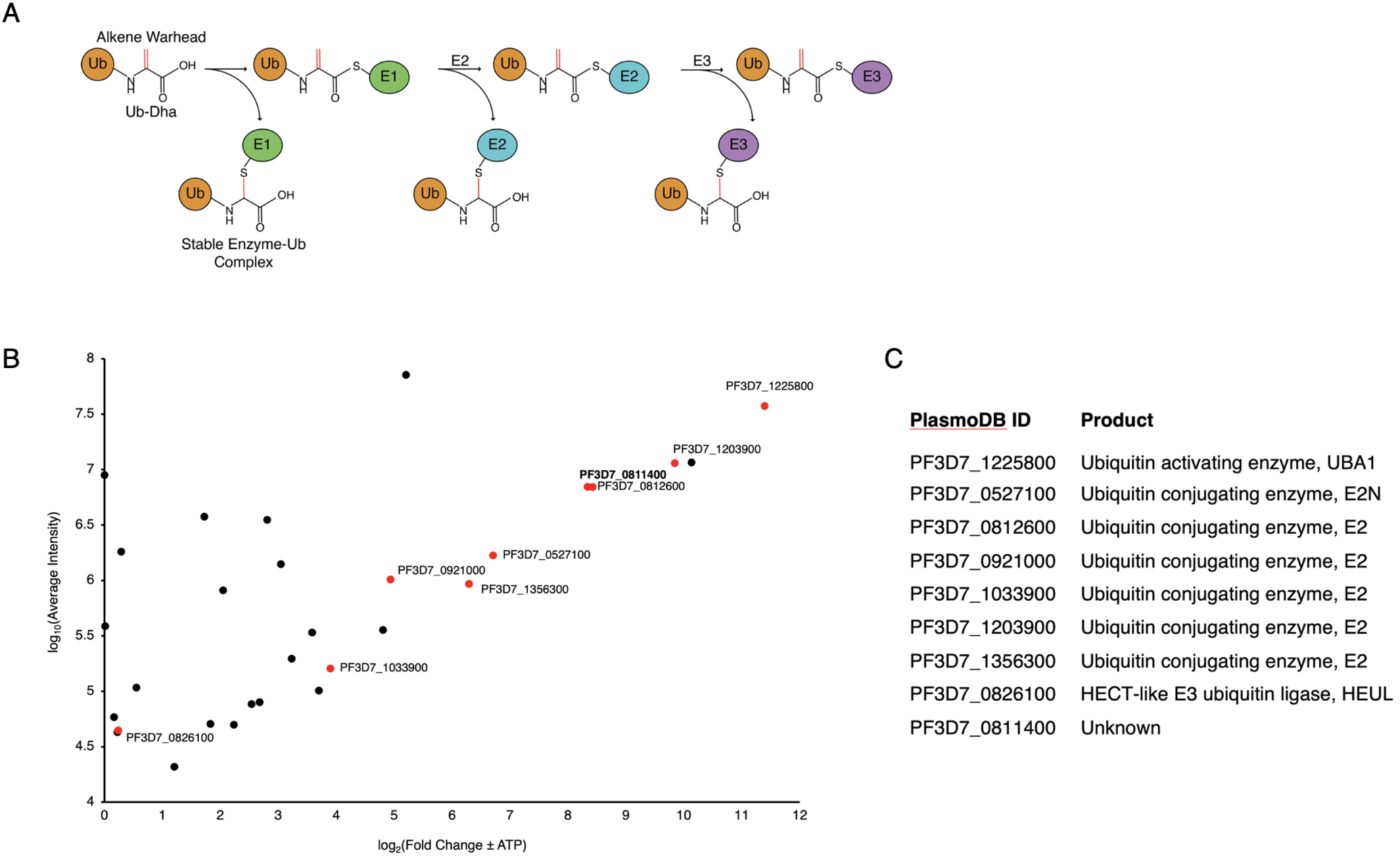
ABP-mediated capture and Identification of *P. falciparum* Ub conjugation machinery. **A)** *Ub-Dha reaction scheme.* Diagrammatic representation of the bipartite reaction mechanism of Ub-Dha within the ubiquitination cascade. Adenylation activates the latent alkene moiety of dehydroalanine as an electrophile through the activity of the E1 ubiquitin activating enzyme. Pathway 1 represents the nucleophilic attack of the alkene moiety of dehydroalanine that results in stable capture of nucleophile-containing, ubiquitin-binding enzymes via thioether bond formation. Pathway 2 represents the native cascade of E1 to E2 to E3 enzyme by subsequent transthioesterification reaction. Pathway 1 can occur at any stage of the ubiquitination cascade. **B)** *P.falciparum proteins captured by Ub-Dha.* Christmas tree plot of biotin-Ub-Dha-trapped proteins from *P.falciparum* lysate identified by LC/MS/MS. The x-axis is fold enrichment and set to start at zero to present enriched hits, and the y-axis is the average iBAQ (LFQ) intensity values. Hits labelled in red are those related to the ubiquitinating enzymes in addition to the unknown PF3D7_0811400. Putative functions of these hits are listed in **C)** *P.falciparum proteins captured by Ub-Dha.* Ubiquitin-related proteins identified by LC-MS/MS using the biotin-Ub-Dha probe. 3 technical repeats were analysed along with the negative control which utilised apyrase to catalyse the conversion of ATP to AMP and inorganic phosphate to effectuate the absence of ATP.

Mixed-stage asexual *Plasmodium falciparum* were saponin-extracted from their host erythrocytes, subjected to freeze-thaw lysis and clarified by centrifugation. The resultant lysate was divided between experimental and control samples. The negative control lysate is depleted of residual ATP by the addition of apyrase which catalyses the sequential hydrolysis of ATP to AMP and inorganic phosphate. This lysate was also not supplemented with ATP during the probe incubation period, unlike the experimental biotin-Ub-Dha reaction. The probe was incubated with control and experiment lysates supplemented with ATP to facilitate probe activation. Proteins of interest, now covalently conjugated to the probe were extracted from cell lysate using neutravidin resin and a series of harsh wash conditions. The eluate from this procedure was submitted for protein identification by mass spectrometry (the mass spectrometry proteomics data have been deposited to the ProteomeXchange Consortium via the PRIDE^20^ partner repository with the dataset identifier PXD056473). Several ubiquitin-related enzymes were identified by the biotin-Ub-Dha probe: one E1 ubiquitin activating and six E2 conjugating enzymes were enriched compared to background (Figure 1B and C). This is in the context of the two known ubiquitin activating enzymes and the 12 predicted ubiquitin conjugating enzymes encoded by *P. falciparum*, although upon inspection, only 10 of these possess putative catalytic cysteines (Supplementary Figure 1). E3 ligases capable of reacting with the electrophilic probes, specifically the four annotated HECT E3 ligases, were not identified above background. However, the HECT E3 ligase PfHEUL was identified by one peptide in one of the technical repeats. A similar number of peptides derived from the Ub-Dha were identified in the ATP supplemented and ATP depleted control experiments, while almost no peptides derived from ubiquitinating enzymes in the control sample. This demonstrated the efficacy of the apyrase treatment in depleting residual ATP and facilitated the reduction of background noise.

### *P. falciparum* E2 conjugating enzymes are active and can undergo autoubiquitination

Several functionally uncharacterised E2 enzymes were identified by activity-based probe; with their putative activity being inferred through their domain homology. PF3D7_0527100, PF3D7_092100, PF3D7_103390, PF3D7_1203900, and PF3D7_1356300 were recombinantly expressed in *E.coli* with an N-terminal His-tag, then purified by immobilised metal affinity chromatography (IMAC) using nickel-affinity resin followed by size exclusion chromatography (SEC). The Plasmodium E1 activating enzyme was also recombinantly expressed in *E.coli* using the codon-optimised plasmid kindly donated by the Holder lab and purified as described in ^3^. A sixth putative E2 enzyme, PF3D7_0812600, was not included in this panel owing to difficulties in PCR amplification.

Assessment of the E1 and E2 enzyme purity was performed using Coomassie-stained SDS-PAGE. For all enzymes expressed and purified, the most prominent band was present at the predicted mass of the representative protein. In the case of PfE1 (PF3D7_1225800), several lower molecular weight bands were observed despite the protein having been exposed to affinity and size exclusion. These bands may represent degradation products (Figure 2A).

**Figure 2.**
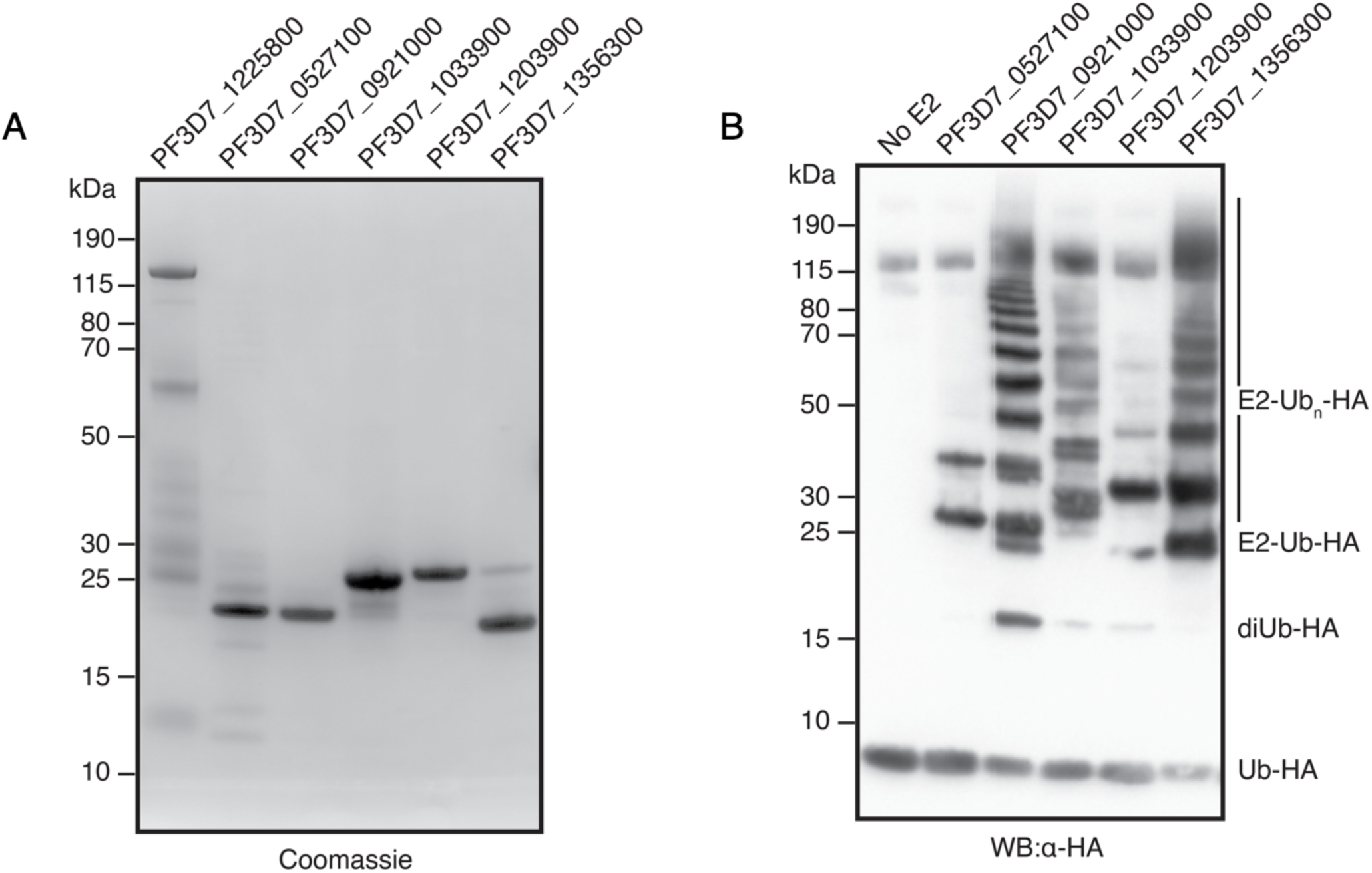
Validation of *P. falciparum* E2 enzyme activity. **A)** *Protein expression and purification.* Coomassie stained SDS-PAGE gel of purified proteins: E1 activating enzyme (PF3D7_1225800), E2 conjugating enzymes (PF3D7_0527100, PF3D7_0921000, PF3D7_1033900, PF3D7_1203900, and PF3D7_1356300). **B)** *In vitro autoubiquitination by P.falciparum E2 enzymes.* Western blot of ubiquitination reaction catalysed by select PfE2 enzymes, visualised by anti-HA antibody. PfE2 enzymes were incubated with ATP, PfE1, and HA-Ub. The products of the reaction were resolved and visualised to reveal signature ubiquitination patterns formed by the unique activity of the different E2 enzymes.

Functional validation of the E2 enzymes was achieved through in vitro ubiquitination assay in the presence of ATP, HA-tagged ubiquitin, and the *Plasmodium* ubiquitin activating enzyme ^3^. The banding pattern resulting from protein conjugation was resolved by SDS-PAGE and visualised by western blotting using an anti-HA antibody (Figure 2B). The presence of a band at the molecular weight of each E2 enzyme in addition to one or more ubiquitins validated their activity.

Three of the PfE2 enzymes tested (PF3D7_092100, PF3D7_1033900, and PF3D7_1356300) exhibited a clear banding pattern consistent with the conjugation of several ubiquitin proteins, in the absence of an E3 ligase. This phenomenon has been observed with characterised human E2 enzymes ^21–23^. This banding pattern can arise through the multi-monoubiquitination of the E2 enzymes, the extension of a monoubiquitinated site to a polyubiquitin chain, or a combination of both. PF3D7_092100, which demonstrated the most distinct ubiquitin conjugate pattern, possesses only 8 lysine residues, and therefore the >10 bands visible at approximate 10 kDa distance must include polyubiquitin chain linkages.

Although, the same is not necessarily true for PF3D7_1033900 and PF3D7_1356300 which contain 14 and 20 lysine residues, respectively. A band consistent with free di-ubiquitin (approx. 17 kDa) was also observed except in the case of PF3D7_0527100. This is suggestive of the formation of a polyubiquitin chain linkage at the catalytic cysteine of the PfE2 enzymes which was subsequently released. Conjugation to PF3D7_0527100 was limited to two bands of sizes amounting to approximately the enzyme plus 1 and 2 ubiquitin modifiers, which may indicate the inability of this enzyme to undergo polyubiquitin chain formation.

### PfHEUL HECT domain functions with a subset of human and P. falciparum E2 enzymes

The essential ligase PfHEUL was also identified as part of the Ub-Dha screen, under the statistically significant threshold, but demonstrating its presence and functionality during the asexual life cycle stage of *P. falciparum*. The sequence of this protein was submitted to bioinformatic analysis by HMMER and BLAST to investigate its function through domain architecture and identification of homologs in model organisms. Owing to the large size of PfHEUL (8591 amino acids, approximately 1 MDa), HMM searches were performed on the N-terminal and C-terminal halves separately. The C-terminal search identified the HECT domain, while the N-terminal search identified a domain of unknown function 908 (DUF908) between residues 176-478. Known DUF908 and HECT domain containing proteins include the homologous HUWE1 E3 ligase (a.k.a. MULE, ARF-BP1, Lasu1, HECTH9) in humans, UPL1/2 in plants, and TOM1 in yeast. Indeed, a BLAST search of PfHEUL against the returned human HUWE1, *A.thaliana* UPL1, and *S.cerevisiae* TOM1 as the top hits. Sequence alignment of these homologs revealed specific conservation of the HECT domain, sharing approximately 50% identity, while upstream there was no discernible sequence similarity (Figure 3A).

**Figure 3.**
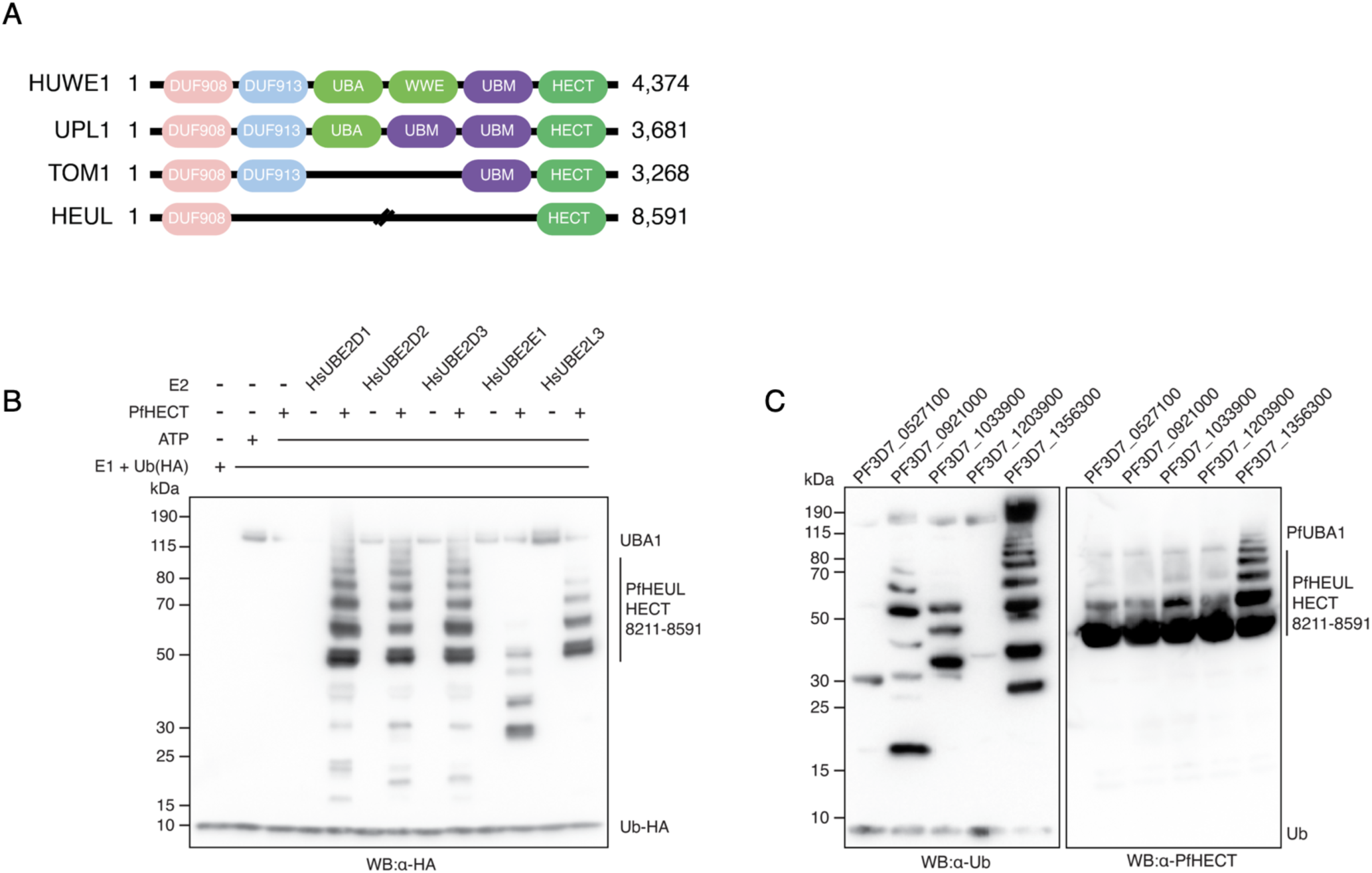
PfHEUL selectivity for E2 enzymes. **A)** *Comparison of PfHEUL to orthologs in model organisms.* Cartoon alignment of PfHEUL with human (Q7Z6Z7), *S.cerevisiae* (Q03280) and *A.thaliana* (Q8GY23) orthologs showing conservation of HECT and DUF908 domains. **B)** *In vitro autoubiquitination of PfHEUL HECT domain using human E2 enzymes.* Anti-HA western blot representing the in vitro autoubiquitination assay used to validate the activity of the PfHEUL HECT domain using human HA-tagged ubiquitin, E1 activating (UBA1), and E2 conjugating enzymes (UBE2D1, UBE2D2, UBE2D3, UBEE21, UBE2L3). All E2 enzymes except UBE2E1 were capable of functioning in concert with PfHEUL HECT. **C)** *In vitro autoubiquitination of PfHEUL HECT domain using P.falciparum E2 enzymes.* Western blots of the in vitro autoubiquitination assay using *P. falciparum* ubiquitination components: E1 activating enzyme (PF3D7_1225800), indicated E2 conjugating enzymes, and the PfHEUL HECT domain. (Left) Western blot using anti-ubiquitin antibodies to identify ubiquitin conjugated proteins following the ubiquitination reaction. (Right) Western blot using mouse derived PfHEUL HECT antibodies to specifically identify PfHEUL HECT in its native and ubiquitinated states.

To validate the activity and thus function of PfHEUL, the HECT domain comprising residues 8211-8591 was tagged with a carboxy-terminal 6His tag and recombinantly expressed in *E. coli* (Supplementary Figure 2). The activity of HECT domains involves the binding of a ubiquitin-loaded E2 conjugating enzyme at the N-lobe of the domain and the subsequent transthiolation reaction at the active site of the C-lobe, where the ubiquitin in transferred from the E2 enzyme onto the ligase cysteine side chain. In vitro and without protein substrate, HECT domains exhibit auto-ubiquitination on their lysine side chains when incubated with upstream components of the ubiquitination cascade^24^. To assess the catalytic function of the HECT domain of PfHEUL an in vitro ubiquitination assay was performed using commercially available human E1 activating, and E2 conjugating components. A panel of five human E2 enzymes was selected based on homology to *Plasmodium* E2 enzymes, characterised promiscuity, and propensity to act as cognate E2 enzymes for other HECT domain proteins. Human E1 and E2 enzymes were mixed with PfHEUL HECT as indicated and incubated at 25 C for 1 hour, in the presence of ATP. Following co-incubation, SDS-PAGE and western blot showed human UBE2D1, UBE2D2, UBE2D3, and UBE2L3 transferred ubiquitin to the PfHEUL (PF3D7_0826100) HECT domain. This was evidenced by banding patterns of approximate 9 kDa spacing which represented the conjugation of multiple ubiquitin proteins onto the HECT domain itself (Figure 3B). UBE2E1 did not exhibit this pattern of HECT domain autoubiquitination demonstrating that this E2 enzyme does not function in concert with the PfHEUL HECT domain.

The panel of *P. falciparum* E2 enzymes (Figure 2) were also tested for PfHEUL compatibility. Both anti-Ub and anti-PfHEUL HECT antibodies were used to distinguish between the ubiquitin-conjugated E2 and E3 enzymes, as autoubiquitination of the E2 enzymes was previously observed. In combination the blots reveal selective compatibility of the Plasmodium enzymes (Figure 3C). Only PF3D7_1356300 was capable of functioning with PfHEUL HECT to bring about polyubiquitination of the HECT domain. The other E2 enzymes only appear to facilitate transfer of a single ubiquitin protein to PfHEUL HECT with progressively diminishing efficiency by PF3D7_1356300, PF3D7_1033900, PF3D7_0527100. PF3D7_0921000 and least of all PF3D7_1203900.

### Identification of C8558 as the active site cysteine of PfHEUL

To further characterise the enzymatic activity of the PfHEUL HECT domain, a C>A mutant was generated (corresponding to C558A in the full length PfHEUL) predicted to correspond to the catalytic cysteine residue of HUWE1 by sequence homology (Figure 4A and B). The expression and purification procedures for this mutant protein were identical to the wild type protein. Circular dichroism (CD) was used to assess the folded state of the mutant protein relative to the wild type. The acquired spectra were averaged across five measurements, and the resultant curve was smoothed and normalised by protein concentration. The CD spectra of both the wild-type PfHEUL HECT domain and the C8558A mutant were almost identical, indicating the mutation did not affect the HECT domain fold (Figure 4C). The spectral minima and maxima also give insight into the secondary structure of the protein. The generated spectra were submitted to the BeStSel deconvolution algorithm (Micsonai et al. 2022) which returned a predicted secondary structure composition of 66% ⍺-helical and 33% β-sheet which is consistent with the mixed secondary structure of helices and sheets in the AlphaFold predicted model (Figure 4A). This result indicates both protein forms are folded, adapt a similar secondary structure, and that the C8558A mutation does not affect the overall secondary structure of the HECT domain.

**Figure 4.**
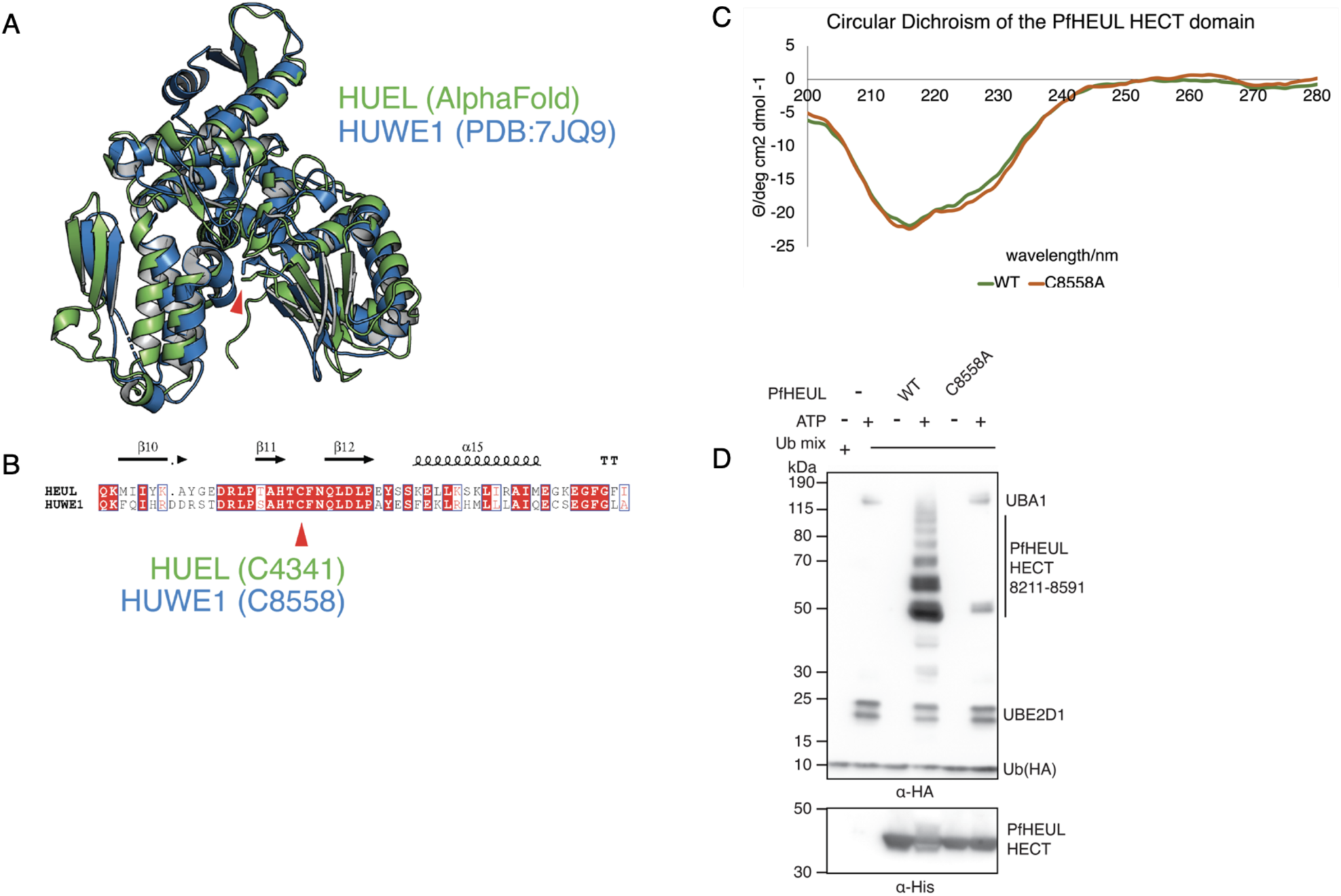
Identification of PfHEUL catalytic cysteine. **A)** *Analysis of PfHEUL HECT domain and catalytic cysteine.* AlphaFold predicted structure of PfHEUL HECT domain (residues 8211-8591) (green) overlaid with crystal structure human HUWE1 (PDB:7JQ9) (blue) and with catalytic cysteines marked by the red arrow . **B)** *Alignment of PfHEUL and human HUWE1 HECT domains.* Rendered in ESPript with catalytic cysteine demarcated by the red arrow. **C)** *Circular Dichroism spectra of wild type (WT) and C8558A mutant PHEUL HECT recombinant protein.* **D)** *In vitro ubiquitination assay.* Anti-HA western blot of an in vitro ubiquitination assay using wild type and C8558A mutant PfHEUL HECT domains to demonstrate the contribution of C8558 to the transthiolation of ubiquitin, and subsequent ubiquitination of lysine side chains.

We next sought to assess the contribution of the putative active site cysteine to transthiolation and ubiquitin transfer using the in vitro assay with human E1 and E2 components. UBE2D1 was used as the cognate E2 conjugating enzyme in this assay as it previously exhibited the clearest ubiquitination banding pattern by western blot in the panel of E2 conjugating enzymes tested. When introduced into the in vitro ubiquitination assay using human E1 and E2 components, the autoubiquitination activity of the C8558A mutant was almost completely abolished compared to the wild type domain (Figure 4D). However, there were 2 bands present at the approximate mass of the HECT domain that are possibly the result of a single transthiolation reaction at a neighbouring non-active site cysteine ^25^.

### Identification Pf3D7_0811400, a novel E2 enzyme

An uncharacterised protein (PF3D7_0811400) was highly enriched relative to background, which suggests this protein is capable of binding ubiquitin or its enzymatic-adducts (e.g. E2 Ub) and possesses a sufficiently reactive cysteine side chain permissive to nucleophilic attack by the Dha moiety. This protein possesses 13 cysteine residues out of a total 608. No functional domains were identified from the primary sequence of this protein by HMMER search, rather it contains a single DUF2009 (Domain of Unknown Function) domain between residues 118-608.

Analysing the predicted 3D structure of Pf3D7_0811400 revealed two distinct domains. The N-terminal domain (amino acids 1-88) includes five short α-helices (h1-h5, Figure 5) and is connected via a 17-amino-acid linker to the C-terminal DUF2009 domain. We performed a 3D structure search of Pf3D7_0811400 using Foldseek^26^ against various databases, including experimental protein crystal structures (PDB) and prediction-based structures (AlphaFold).

**Figure 5.**
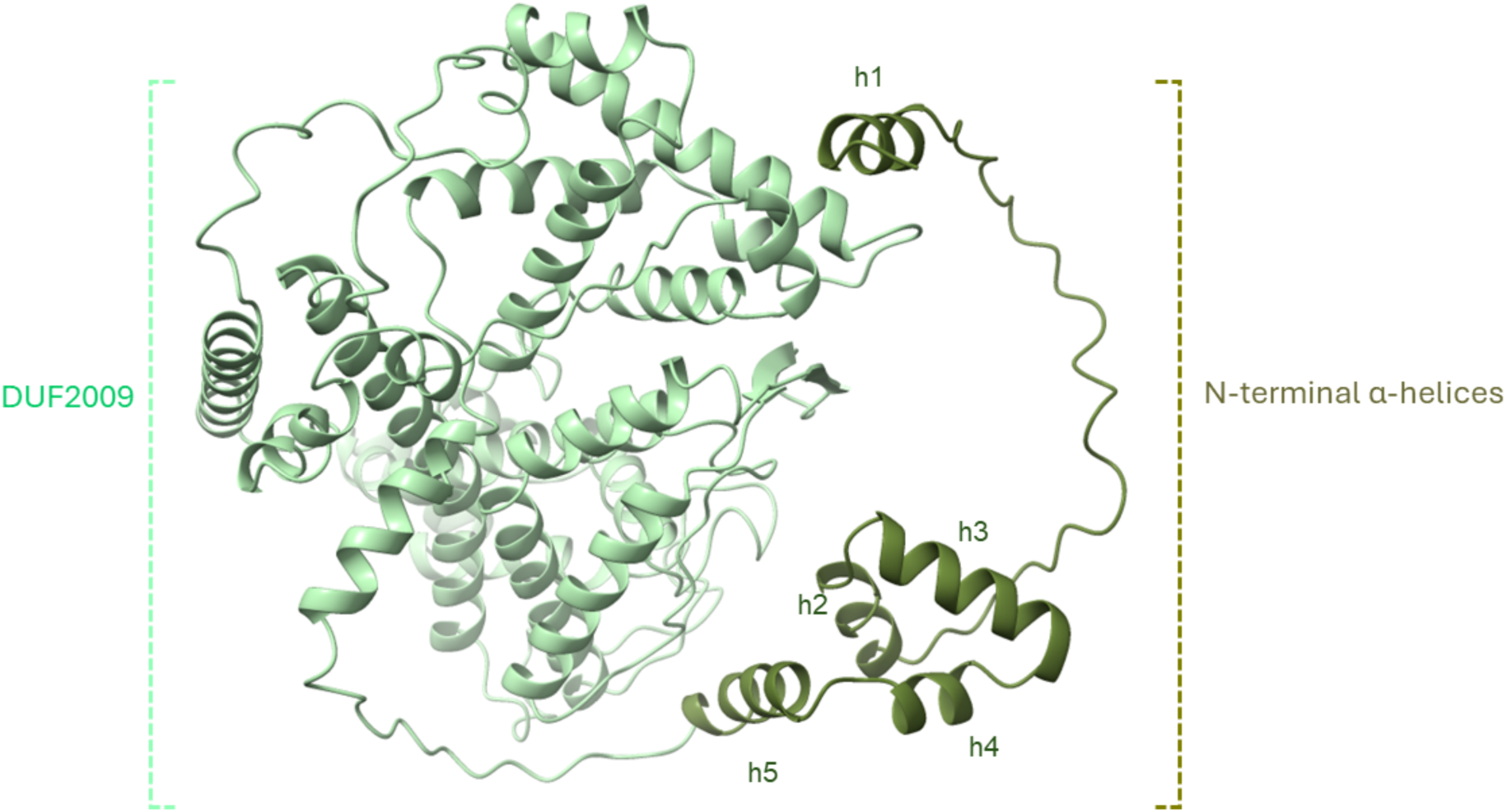
AlphaFold-predicted structure of Pf3D7_0811400. The N-terminal α-helices (coils) are coloured dark green, while the C-terminal DUF2009 domain is shown in sea green.

The structure matched several uncharacterized proteins from other organisms, specifically putative adenylosuccinate lyase (ADSL) in *Toxoplasma gondii* and an unidentified B-box containing protein in a few organisms such as *Synchytrium microbalum* and *Rhizophagus irregularis* (Supplementary Table 1). Since many E3 ligases contain B-box domains, we examined this further.

A typical B-box domain contains histidine and cysteine residues that coordinate with zinc. Two types of B-boxes exist: B-box1, which spans 50–60 amino acids with a zinc-binding consensus sequence of C-X₂-C-X₇–₁₂C-X₂-C-X₄-C-X₂-[C/H]-X₃–₄-H-X₄–₉-H [C₅(C/H)H₂], and B-box2, which spans 35–45 amino acids with a consensus sequence of C-X₂-H-X₇–₉-C-X₂-[C/D/E]-X₄-C-X₂-C-X₃–₆-H-X₂–₄-[C/H] [CHC(C/D/E)C₂H(C/H)] ^27^. Although the DUF2009 domain of Pf3D7_0811400 is structurally conserved, it does not possess the typical B-box structure or signature motifs (Supplementary Figure 3).

Pf3D7_0811400 contains 13 cysteine residues within its 608-amino-acid sequence. None of these cysteines are located in the N-terminal region, which led us to hypothesise that the N-terminus may be involved in protein-protein interactions, potentially with E3 ligases or E3 complex components, while the C-terminus interacts with ubiquitin. Conservation analysis was conducted using a ConSurf model, which estimates the evolutionary conservation of amino acid positions based on phylogenetic relationships among homologous sequences^28^. This analysis revealed that the highest conservation levels are found in the C-terminus of Pf3D7_0811400, particularly around two cysteine residues, C572 and C584. C572 is 100% conserved across all homologs, while C584 shows 95% conservation (Supplementary Figure 4). Structural analysis indicated that the thiol side chain of C572 is located in a loop at the base of a conserved and accessible pocket, whereas the side chain of C584 is buried between two α-helices. Protein-protein interaction modelling using AlphaFold suggested that ubiquitin interacts with C572 of Pf3D7_0811400 (Figure 6).

**Figure 6.**
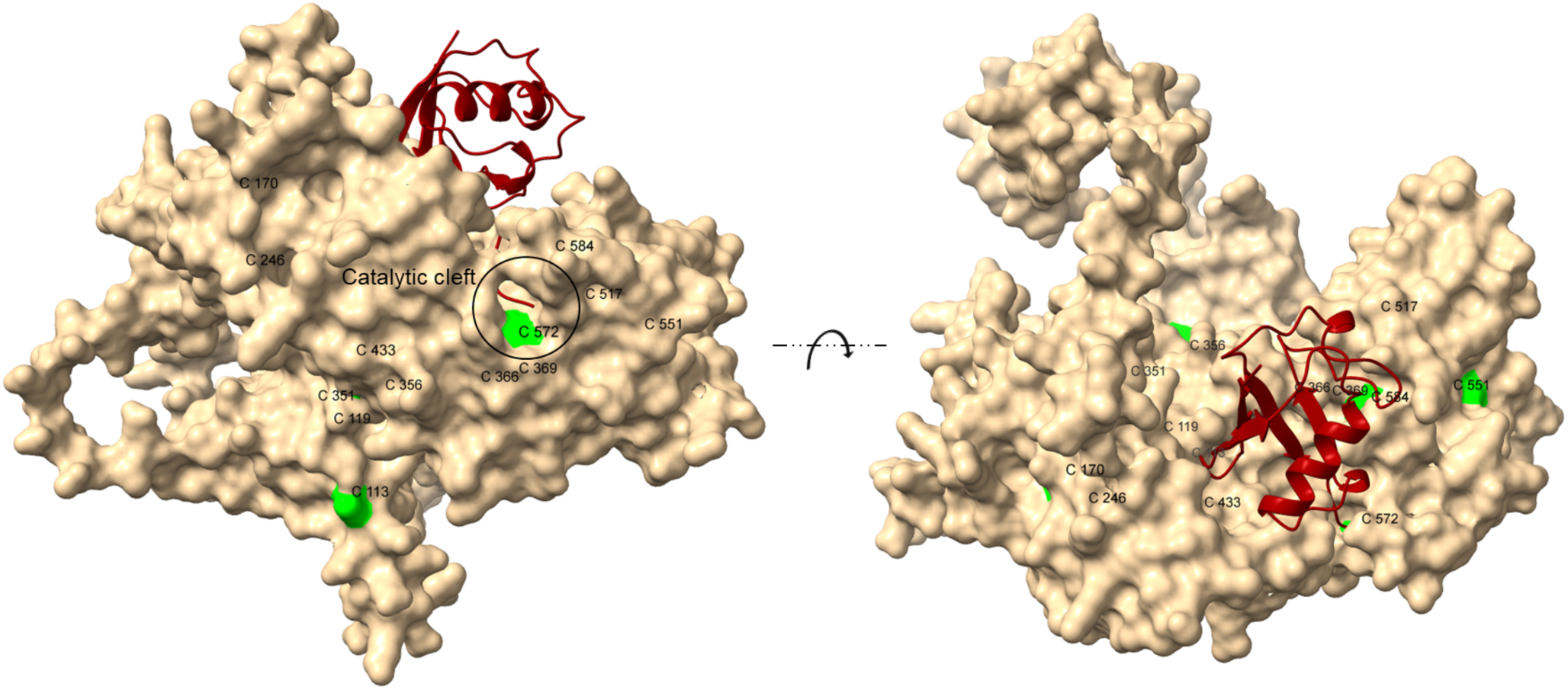
AlphaFold-predicted interaction between Pf3D7_0811400 (beige) and ubiquitin (red). All 13 cysteine residues in Pf3D7_0811400 are labelled and highlighted in green. The C-terminal glycine (Gly76) of ubiquitin interacts with C572, which acts as the catalytic cysteine in the core of the catalytic cleft.

### The N-terminus of Pf3D7_0811400 is critical for interacting with PfRbx1

Despite its dissimilarity to other E2 and E3 ubiquitin enzymes, Pf3D7_0811400 was a particularly robust hit in the mass spectrometry screen. As such, we suspected it to be an E2 Ub conjugating enzyme and set out to test this hypothesis and to understand which domain is essential for its activity. To start with, the five N-terminal α-helices (amino acids 1–88), as predicted by InterPro, were truncated to investigate their role in protein-protein interactions (Figure 7A). HEK293 cells were transfected with either an empty vector, FLAG-tagged Pf3D7_0811400, or FLAG-tagged Pf3D7_0811400ΔN, a N-terminal truncation missing the first 88 amino acids. Whole cell lysates were subjected to probe labelling by adding biotin-Ub-Dha in the presence of Mg-ATP and human E1 enzyme. The proteins were then co-immunopurified using anti-FLAG magnetic resin and eluted with Laemmli buffer. Western blot analysis with HRP-conjugated streptavidin showed that the biotinylated probe co-immunopurified with both the wild-type and truncated proteins (Figure 7B, top panel).

**Figure 7.**
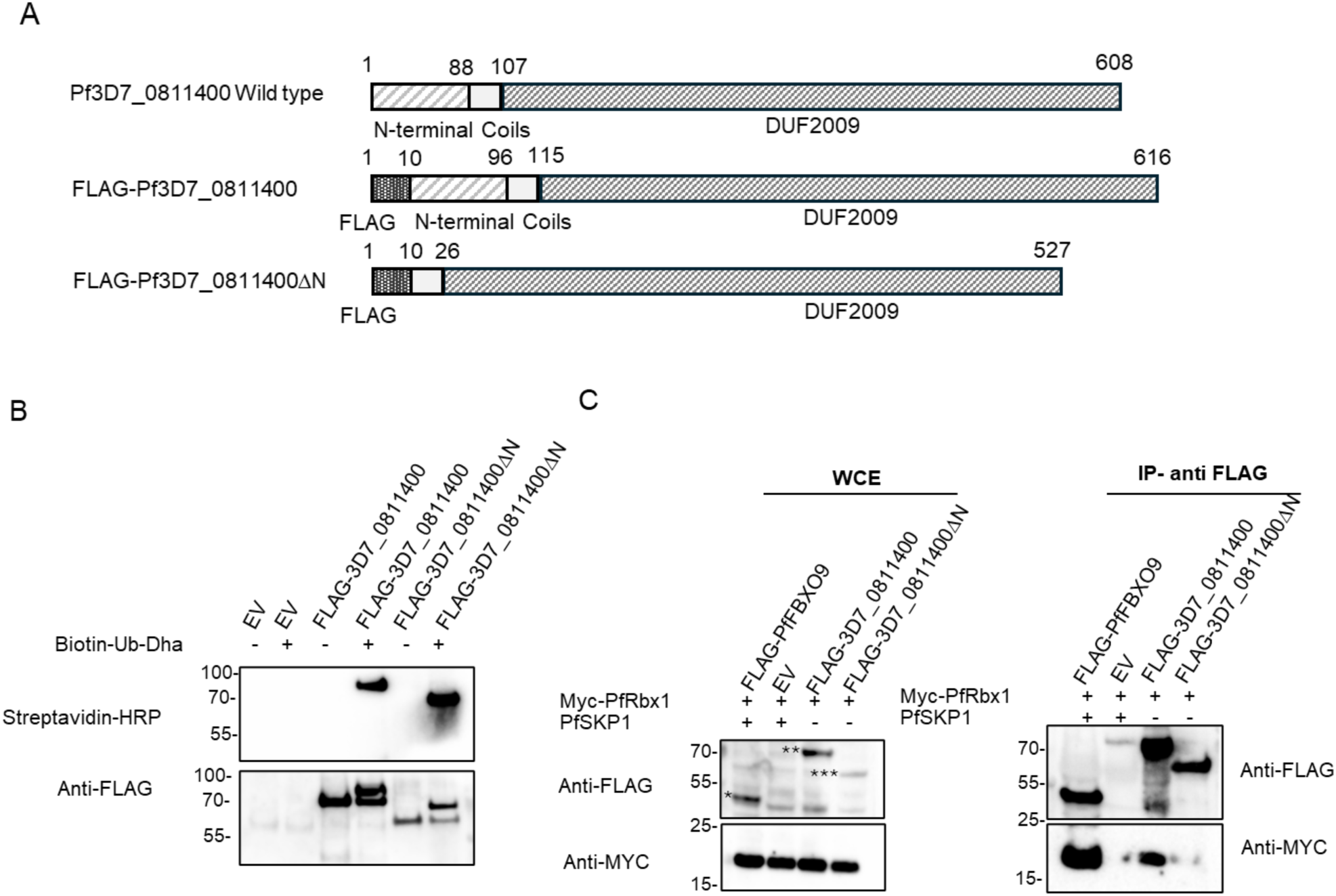
Pf3D7_0811400 interaction with Ub-Dha probe and PfRbx1. **A)** *Schematic of Pf3D7_0811400.* Domains and truncations used to generate protein for labelling and interaction studies are shown. Five predicted N-terminal α-helices (amino acids 1–88) were truncated to investigate their role in protein-protein interactions**. B)** *Validation of Pf3D7_0811400 interaction with Ub-Dha probe.* HEK293T cells were transfected with either an empty vector, FLAG-tagged Pf3D7_0811400, or FLAG-tagged Pf3D7_0811400ΔN. Whole cell lysates were labelled with biotin-Ub-Dha in the presence of Mg-ATP and additional E1 enzyme. The proteins were then co-immunopurified on anti-FLAG resin. Proteins were visualised by immunoblot with HRP-conjugated streptavidin (top panel) or Anti-FLAG (bottom panel) to confirm a ∼10 kDa shift in protein size in the presence of the probe. **C)** *Pf3D7_0811400 interaction with PfRbx1 via its N-terminal domain*. MYC-tagged PfRbx1 was co-expressed with either FLAG-tagged Pf3D7_0811400 WT (indicated by**) or the ΔN truncated mutant (indicated by***) in HEK293T cells. FLAG-tagged PfFBXO9 (indicated by*) in the presence of PfSkp1 was used as a positive control, while an empty vector (EV) in the presence of both PfRbx1 and PfSkp1 served as a negative control. Immunopurification using anti-FLAG beads was performed on all samples and proteins were visualised by anti-MYC or anti-FLAG. WCE=whole cell extract. Images of full membranes are included in **Supplementary** Figure 5.

Anti-FLAG immunoblotting confirmed an approximate 10 kDa shift in protein size in the presence of the probe (Figure 7B, bottom panel), indicating that the N-terminal domain is not involved in the interaction with ubiquitin.

Since Pf3D7_0811400 was also captured in a PfCullin-2 pulldown we had previously performed^29^ and since many E2 enzymes interact with Rbx1 as part of Cullin RING ligase complexes, we set out to test this interaction. MYC-tagged PfRbx1 was co-expressed with either FLAG-tagged full-length Pf3D7_0811400 or the N-terminal truncation. PfFBXO9 in the presence of PfSkp1 or empty vector were included as positive and negative controls, respectively. Anti-FLAG beads were used to immunopurify all samples and proteins were visualised by anti-FLAG or anti-MYC immunoblot. As shown in Figure 7C, Pf3D7_0811400 does co-precipitate with PfRbx1, and its N-terminus appears to be critical to mediating this interaction, further supporting its characterisation as an E2.

### C-terminal end of Pf3D7_0811400 mediates binding to ubiquitin

Since the N-terminus of Pf3D7_0811400 was found not to be essential for its interaction with the Ub-Dha probe, we next focused on its C-terminus which contains the DUF2009 domain and the putative catalytic cysteine C572 as discussed above. We mutated C572 to an arginine and repeated the probe-interaction experiment in the presence or absence of ATP. While the wild-type full-length protein bound the probe robustly, the C572R mutation entirely abrogated this interaction as demonstrated by both streptavidin and anti-FLAG immunoblot (Figure 8A). To confirm that this interaction is indeed mediated by a direct binding to ubiquitin via the C572R cysteine residue, we repeated the assay but this time used HA-Ub in the place of biotin-Ub-Dha probe. Again, in the presence of ATP, wild-type Pf3D7_0811400 was able to bind ubiquitin whereas the cysteine mutant enzyme did not (Figure 8B, left panel). Notably, the reaction with the wild-type protein did not result in a ladder as seen in the positive control reactions we ran in parallel using the *P. falciparum* Pf3D7_0921000 E2 and the human E2, UB2R1 (Figure 8B, right panel). Notably, while all identified E2 enzymes share a conserved catalytic HPN motif in addition to the catalytic cysteine, this motif is absent in Pf3D7_0811400, suggesting a different mechanism of action.

**Figure 8.**
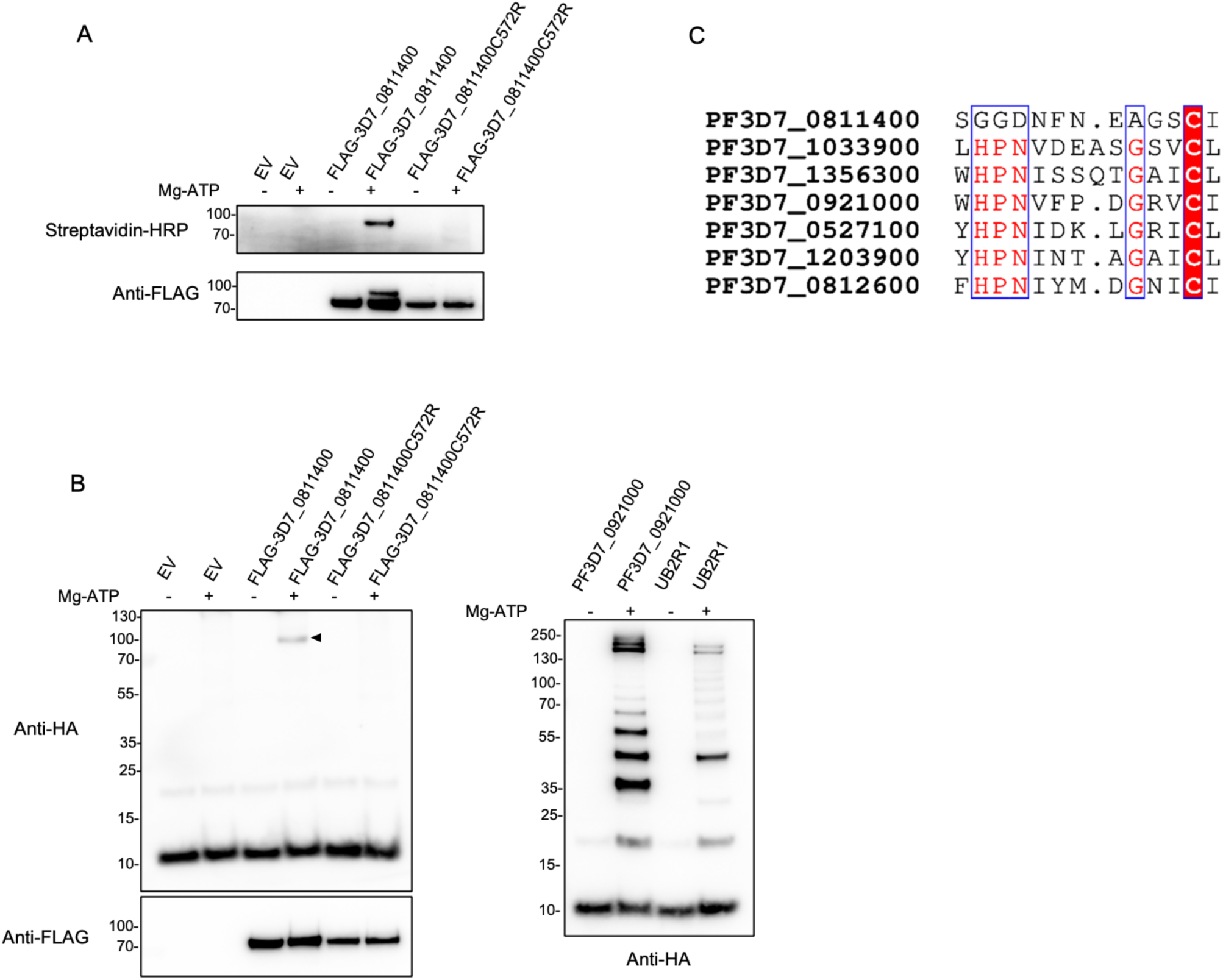
Identification of PF3D7_0811400 as an E2 and C572 as its catalytic cysteine. **A)** *Biotin-Ub-Dha-reactivity of wild type and C572R proteins.* Wild-type and C572R mutant proteins were expressed in HEK293T cells, purified using anti-FLAG beads and eluted by incubation with a 3XFLAG peptide solution. Purified proteins were subjected to probe labelling in the presence or absence of Mg-ATP. (Images of full membranes are included in **Supplementary** Figure 6). **B)** *Ubiquitin-HA-reactivity of wild type and C572R proteins.* Purified FLAG-tagged Pf3D7_0811400 or the C572R mutant was used in an in vitro auto-ubiquitination assay. HA-tagged ubiquitin was added to allow detection by anti-HA immunoblot (left panel). The arrow highlights the weak and transient, but highly specific, formation of a thioester-linked E2 ubiquitin (E2∼Ub) conjugate in the wild-type E2. PF3D7_0921000, a *Plasmodium* E2 characterised earlier in this paper, and UB2R1, a human E2, were used as positive controls (right label). **C)** *Comparison of Pf3D7_0811400 with other P. falciparum E2 enzymes.* Sequence alignment showing the conserved catalytic HPN motif in all of the described Pf E2 enzymes but not in Pf3D7_0811400.

## Discussion

The ubiquitin landscape of *Plasmodium* has been predicted in silico but biochemical exploration is limited ^5,29–33^. A profile of the ubiquitin pathway acting at the pathology-causing stage of *P. falciparum* is a useful and important platform from which to initiate programmes of study and screening using ubiquitin enzymes identified as viable therapeutic targets. Previous work has demonstrated the functional activity of the E1 activating enzyme in vitro and in vivo^3^ and the E2 enzyme PF3D7_1203900 in vitro^30^. The E2 enzyme PF3D7_0527100 was also shown to be active in vitro and negatively regulated through S106 phosphorylation by PfPK9^34^, with subsequent work demonstrating its involvement in K63 polyubiquitin linkage formation in vivo^35^. Several structures of Plasmodium E2 ubiquitination enzymes have been solved as part of larger structural elucidation programmes^36^, including: PF3D7_0527100 (2R0J) in complex with PF3D7_0305700 (2Q0V; 3E95), PF3D7_1033900 (2ONU), PF3D7_1345500 (2H2Y) and the *P.vivax* homologue (2FO3). Together, the structural data and the functional validation presented in this study provides a basis for structure-guided drug design.

One putative E3 ligase (PF3D7_0826100 PfHEUL) was identified using ABPs in this study, albeit with low confidence, although its presence during asexual development is supported by transcriptomic data^37^. There may be other factors limiting the probes’ reactivity to these enzymes and their subsequent detection. Protein levels may be transient and low abundance making peptide detection by mass spectrometry challenging. The cascading mechanism of Ub-Dha leads to attrition of the ubiquitin machinery following capture of enzymes at each stage. Thus, the capture of E3 ligase may happen at levels below the detection limit of this approach.

There are a total of 4 HECT domain-containing E3 ligases annotated in the *Plasmodium falciparum* genome. Of the four, only PfHEUL was found to be essential by piggyBac mutagenesis in *P. falciparum* ^38^. Based on the structure of distant homologs in humans and fungi, PfHEUL may adopt a large solenoid structure where the HECT domain catalyses ubiquitin ligation to specific substrates recruited by the extensive protein binding modules upstream of the catalytic domain^39,40^. Mutations in this enzyme have been linked to pyrimethamine resistance, however the mechanism has not been investigated. The position of these mutations were found upstream of the HECT domain and likely alter protein recruitment as opposed to ubiquitin transfer^41,42^. This class of E3 ubiquitin ligases has also proven to be a tractable and specific targets for small molecule inhibition, with several sites of inhibition identified across the HECT domain^43,44^. Together, these factors suggested PfHEUL would be a suitable target for antimalarial therapeutics.

PfHEUL is one of the largest proteins in the *P.falciparum* proteome. At 1 MDa, direct study of the protein in its entirety is problematic owing to issues arising in cloning and recombinant protein production. To circumvent these obstacles the catalytic 45 kDa HECT domain was examined in isolation. The sequence of the HECT domain of this protein is highly conserved across *Plasmodium* species, indicating the functional importance this catalytic domain plays within this E3 ligase and across *Plasmodium* biology. The protein sequence upstream of the HECT domain is less conserved, which suggests species-specific variation with regards to substrate recruitment.

Within the activity assay, all tested E2 enzymes were capable of conjugating ubiquitin, although unique polyubiquitination patterns were observed most clearly with PF3D7_092100, PF3D7_103390, and PF3D7_1356300. E2 enzymes have been demonstrated to catalyse auto(poly)ubiquitination in the absence of E3 enzymes, however the physiological relevance of this is unclear^23^. It may represent the ability of these E2 enzymes to form polyubiquitin chains at their active site which they could transfer en masse in vivo, as opposed to a model where monoubiquitination of a substrate lysine primes the protein for successive ubiquitin chain extension by single ubiquitin proteins.

PF3D7_0527100 (PfUBC13) and its homologs are associated with K63 polyubiquitination, although no extensive polyubiquitination pattern was produced by this enzyme in the in vitro assay^35^. There are ubiquitin-conjugating enzyme variants (UEVs) that lack the active site cysteine and are thus non-catalytic; homologs of PfUBC13 have been found to associate with UEVs to modulate their E2 activity and linkage specificity^45^. There are two examples of these UEVs in *Plasmodium*: PF3D7_0305700 (PfMMS13) and PF3D7_1243700. As mentioned, the co-structure of PfUBC13:PfMMS2 has been solved, demonstrating a stable interaction. The formation of this complex may be required for ubiquitin polymerisation (via K63).

Studies of the *Plasmodium falciparum* protein interactome have shown physical association of PfMMS2 with several E2 enzymes: ubiquitin-conjugating enzymes PF3D7_0527100, PF3D7_092100, PF3D7_1203900, PF3D7_1356300, as well as SUMO-conjugating enzyme PF3D7_0915100, and NEDD8-conjugating enzyme PF3D7_1245300^46^. Future studies of the utility of these accessory E2 enzymes in concert with catalytic E2 enzymes may reveal functional modulation relative to the E2 enzyme alone, expanding the *Plasmodium* ubiquitin and ubiquitin-like small modifier landscape.

The E2 superfamily, classified into 17 families in eukaryotes, typically shares a conserved 150-amino-acid core known as the Ubiquitin Conjugation (UBC) domain, which includes a catalytic cysteine essential for thioester bond formation with ubiquitin. In catalytically active E2s, the active site generally contains an acidic residue, usually aspartate, or a phosphorylatable serine, known as the conserved E2 serine/aspartate (CES/D) site. Together with the catalytic cysteine and the His-Pro-Asn (HPN) motif, this CES/D motif forms a defining fingerprint of the E2 superfamily, crucial to their role in the ubiquitination cascade^47^. For instance, in the human E2, Ube2I, key residues such as Asn85 of the HPN motif, along with Tyr87 and Asp127, are essential for lowering the pKa of substrate lysine, facilitating its nucleophilic attack on the E2Ub thioester bond^48^ . While active E2s are regulated by phosphorylation at their catalytic site, inactive members, like UEVs, lack this conserved catalytic cysteine. However, families 5 and 13, classified as Noncanonical Ubiquitin Conjugating Enzymes (NCUBE), show unique structural reorganisations around the catalytic cleft and do not consistently maintain the HPN motif^49^.

In our study, we identified a novel active E2 enzyme that lacks the HPN motif, the CES/D site, and other typical E2 features, making it unlike any E2 characterised to date. With a molecular weight of ∼71 kDa and a DUF2009 domain, this enzyme may represent a new NCUBE family member. Although the presence of a catalytic cysteine has been reported in a few DUF containing proteins such as DUF62^50^, its presence within the DUF2009 domain is unprecedented and could provide new insights into E2 enzyme function, suggesting that similar E2s may exist in other organisms. This discovery also marks the first identification of an entire active site within a DUF2009 domain, opening avenues for further study of this protein family. From an evolutionary perspective, the high similarity of Pf3D7_0811400’s DUF2009 domain in many B-box proteins, particularly in certain fungal parasites, suggests that the active site within this domain has been evolutionarily conserved. In contrast, the protein-protein interaction site at the N-terminus appears to have undergone significant evolutionary shifts. It has likely transitioned from a metal-binding B-box domain to coiled helices as an interaction interface, adapting the protein’s function to meet the specific requirements of the parasite under natural selection pressures. This highlights the evolutionary flexibility of E2 enzymes, enabling them to respond to environmental and functional demands.

The results detailed in this study demonstrate the applicability of ubiquitin probes for the identification and study of (de)ubiquitination machinery in *Plasmodium*. The validation of several ubiquitin E2 enzymes and their capacity for autoubiquitination has given insight into the machinery of the *Plasmodium* ubiquitin pathway and provided the foundation for the characterisation of novel antimalarial drug targets.

## Acknowledgements

We thank Dr. Andrew Blagborough for helping generate the anti-PfHEUL antibody. We are very grateful for expert help with mass spectrometry analysis by the Discovery Proteomics Facility led by Iolanda Vendrell and Roman Fischer (University of Oxford). This work was supported by a Wellcome Trust Doctoral Training Program studentship held by CS and a MRC grant (MR/W025566/1) held by KAT. APF and BMK were supported by the Chinese Academy of Medical Sciences (CAMS) Innovation Fund for Medical Science (CIFMS), China [grant number: 2018-I2M-2-002].

## Author Contributions

CS and MH generated all the data and figures and KAT designed and directed the research. GJHN contributed reagents and protocols for the ABP experiments. APF and BMK performed the proteomics analysis. CS, MH and KAT wrote the manuscript with input from GJHN, ABP and BMK.

