## Supplemental Figures for "Identification of a new *Plasmodium falciparum* E2 ubiquitin conjugating enzyme"

**Supplementary Figures:**

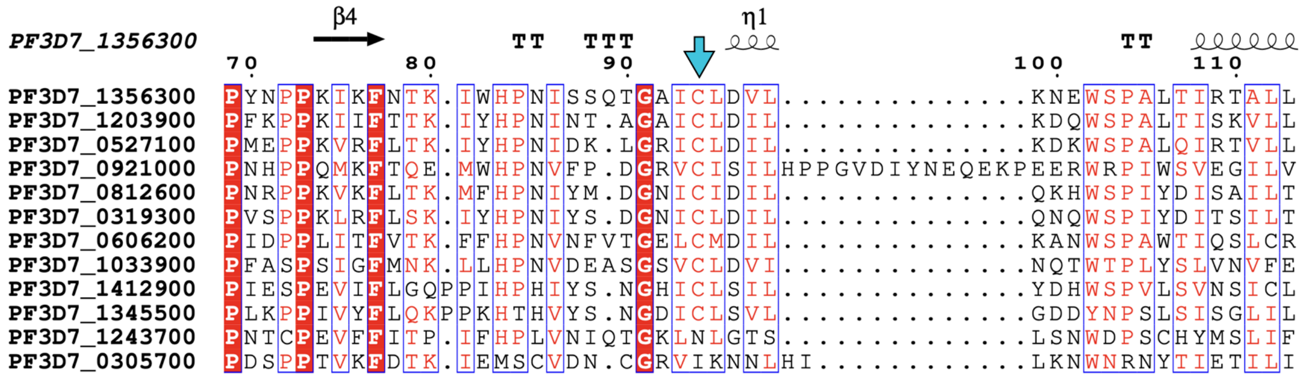

**Supplementary Figure 1. Alignment of P. falciparum putative E2 enzymes.** MUSCLE sequence alignment of PfE2 enzymes rendered in ESPript. All annotated PfE2 enzymes contain a conserved cysteine residue (blue arrow), here at numbered residue 94, except two putative UEV enzymes PF3D7_1243700 and PF3D7_0305700.

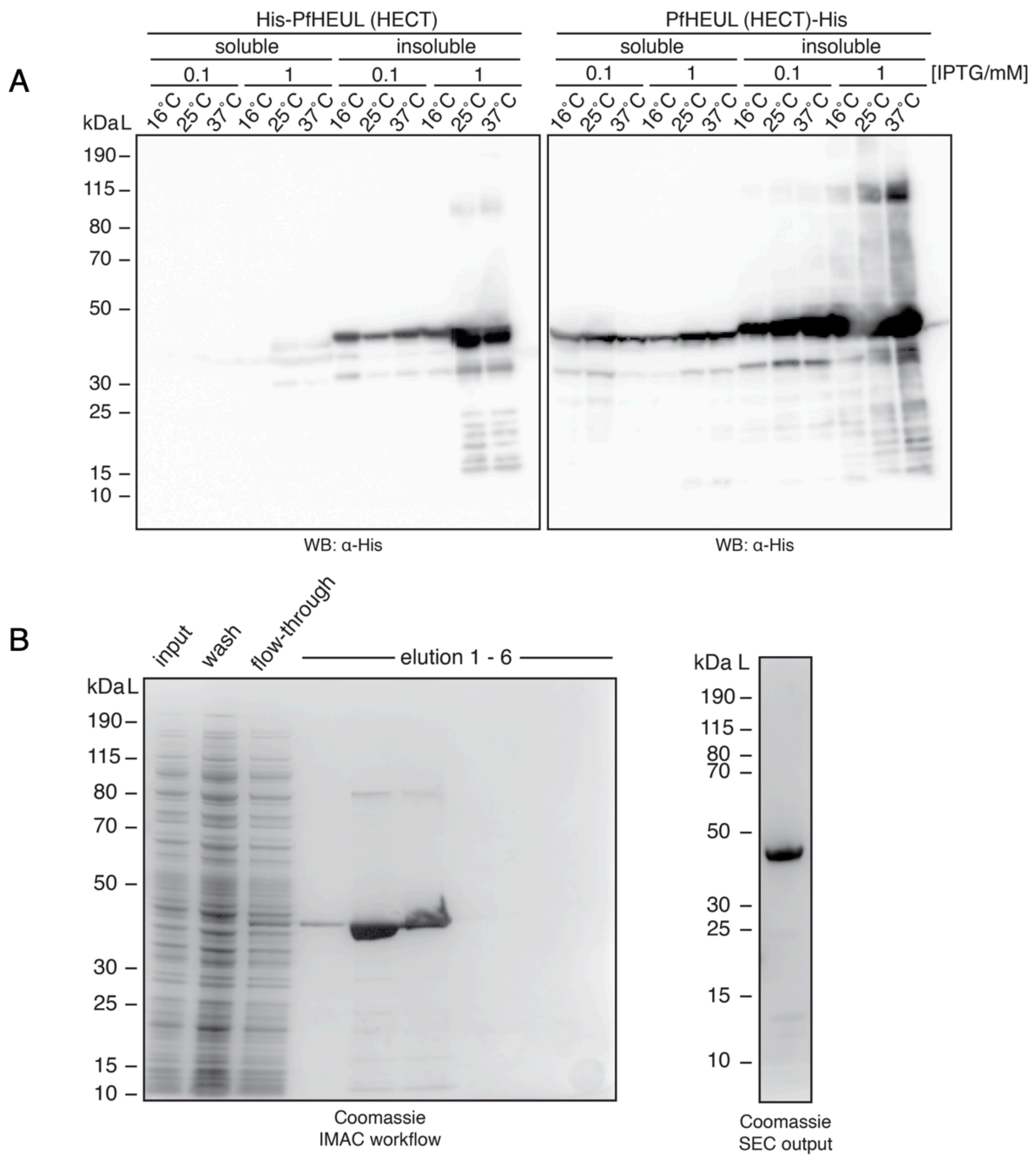


**Supplementary Figure 2. Expression and purification of PfHEUL HECT domain. A)** *Western blot of His-tag PfHEUL HECT (~45 kDa) expression trial under varying incubation temperature and IPTG concentration.* Bacteria were chemically lysed and the soluble and insoluble fractions separated by centrifugation prior to resuspension and SDS-PAGE. **B)** *Nickel affinity and desalting workflow assessing input lysate, wash steps, and elution.* PfHEUL was eluted maximally by the third elution fraction, and these fractions were pooled and subjected to SEC. This sample demonstrated a single band at the expected molecular weight of PfHEUL.

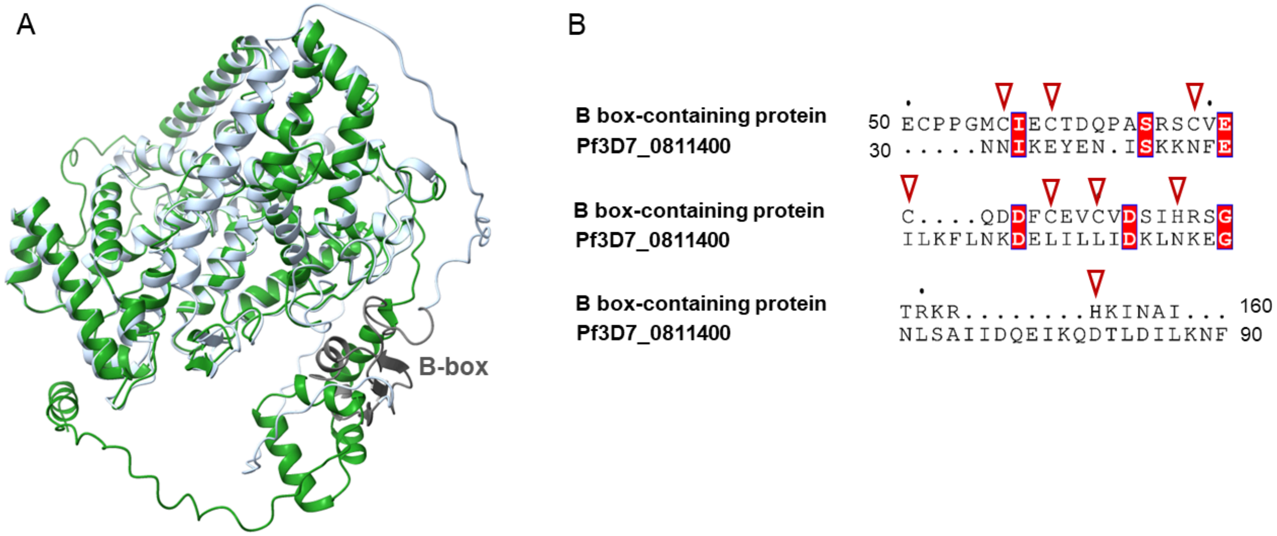


**Supplementary Figure 3. A.** AlphaFold-predicted 3D-structure alignment of Pf3D7_0811400 (green) and *Sphaeroforma arctica* B box-type domain-containing protein (silver). While the C-terminal DUF2009 domain shows a high level of similarity between the two, the B-box domain (highlighted in dark grey) is absent in the N-terminus of Pf3D7_0811400. **B.** Amino acid sequence alignment of the B-box domain from the B box-type domain-containing protein with the corresponding N-terminal region of Pf3D7_0811400. Cysteines and histidines involved in zinc interaction, indicated by red arrows, are present in the B-box protein but absent in Pf3D7_0811400.

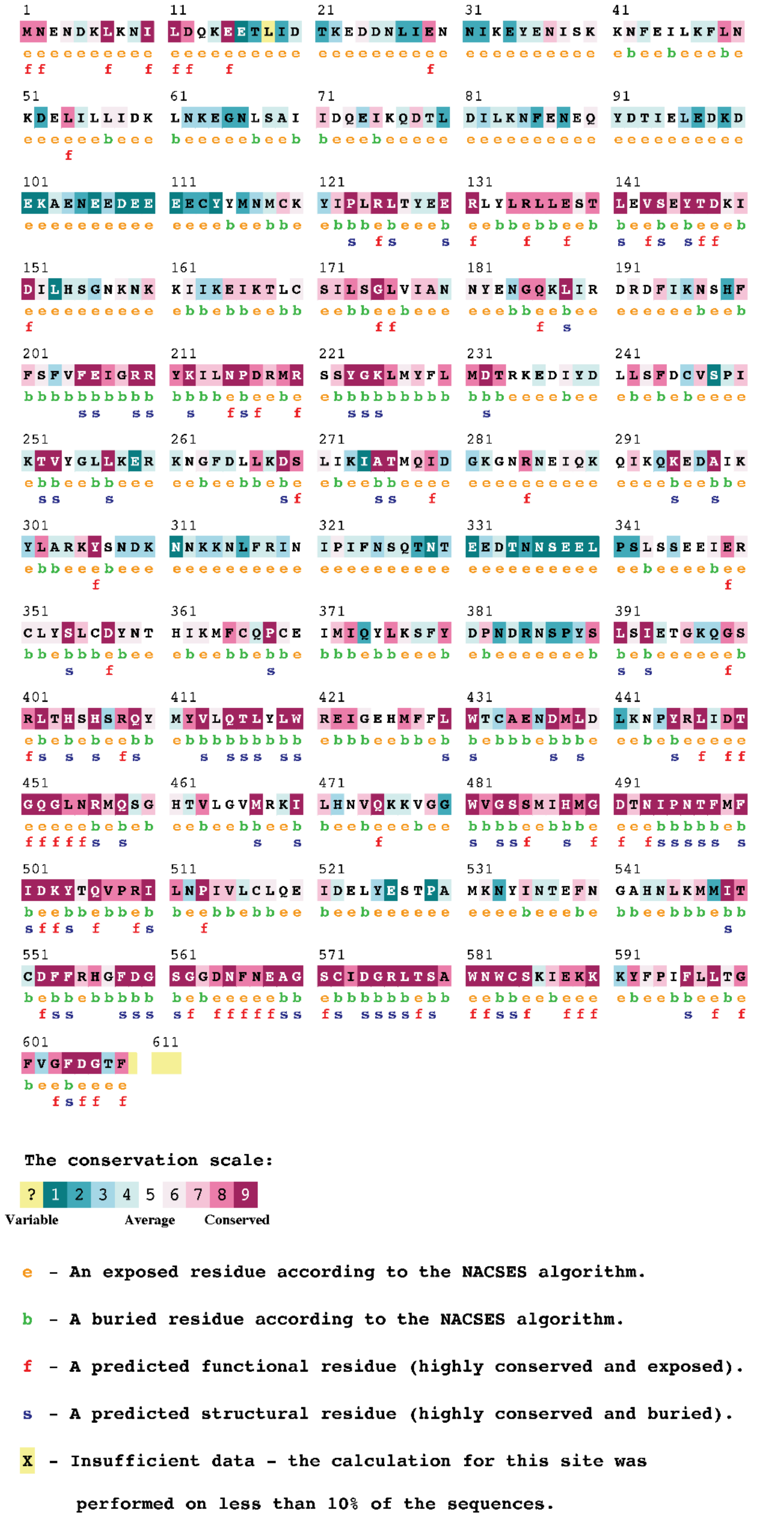


**Supplementary Figure 4. Consurf output for Pf3D7_0811400.** The Consurf web server[19] was used to identify orthologs based on the protein sequence of Pf3D7_0811400. The output was used to colour-code the input PDB structural file (accessed through the AlphaFold web server) based on the computed conservation score for each amino acid residue. Residues are colour-rendered according to the conservation index, with annotations corresponding to the key.

A

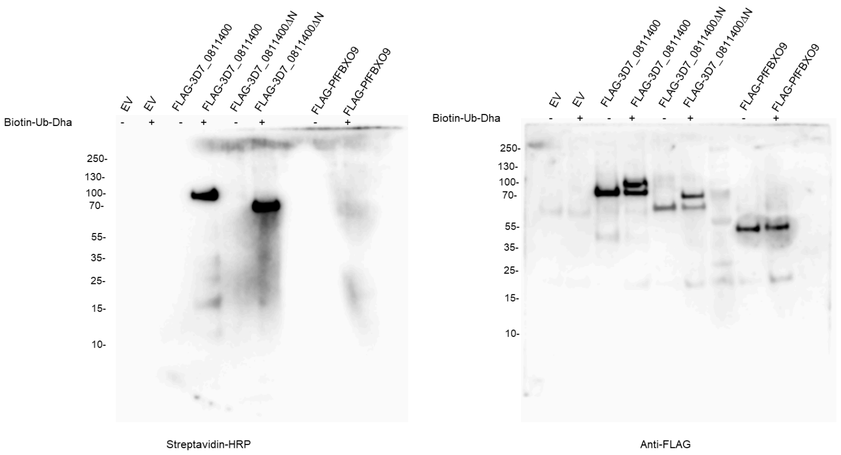


B

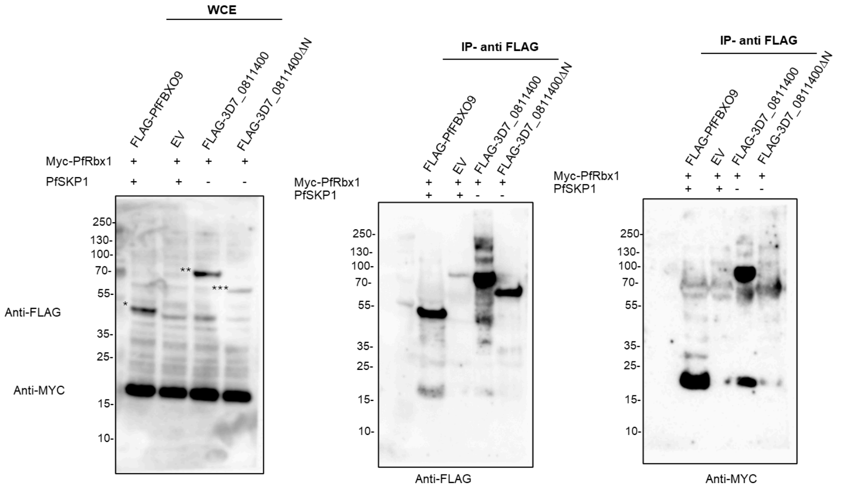


**Supplementary Figure 5. Full membranes for Figure 7 immunoblots.**
**A)** Full membranes for Figure 7B streptavidin-HRP (left) and anti-FLAG (right) immunoblots. FLAG-tagged PfFBXO9 served as an additional negative control. **B)** Full membranes for Figure 7C. The whole cell extract (WCE) membrane was sequentially probed with both anti-FLAG and anti-MYC antibodies. The immunopurified proteins were also probed with both antibodies, but on separate membranes (middle image is anti-FLAG and right image is anti-MYC). FLAG-tagged Pf3D7_0811400 WT (indicated by**); ΔN N-terminal truncated mutant (indicated by***); FLAG-tagged PfFBXO9 (indicated by*).

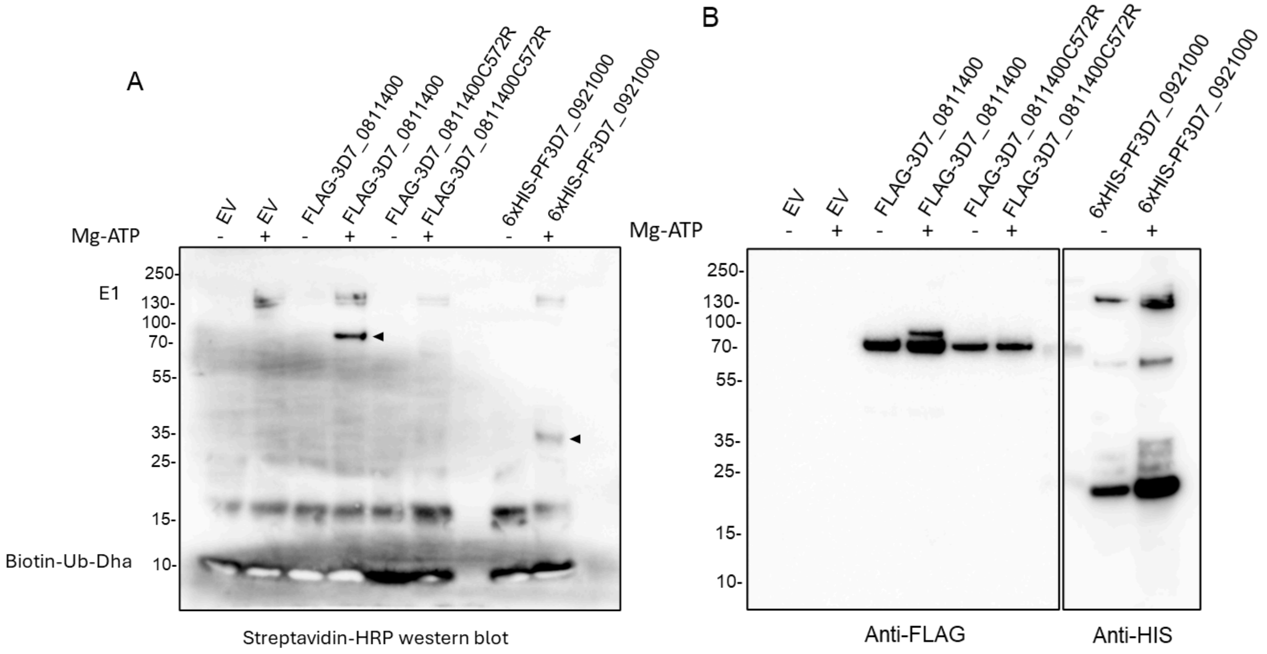


**Supplementary Figure 6. Full membranes for Figure 8A.**

Panels A and B show the full membrane for western blots shown in Figure 8A. Here, a positive control is also included, PF3D7_0921000, a *P. falciparum* E2 enzyme characterised earlier in this paper. Arrows indicate the signals corresponding to probe interactions with each E2. The FLAG and HIS blots represent the loading of PF3D7_0811400 and PF3D7_0921000, respectively.

**Sup table 1:**

PF3D7_0811400 protein structure alignments against large protein structure collections by Foldseek Server.
